# A dual voltage clamp technique to study gap junction hemichannels in astrocytes cultured from neonatal rodent spinal cords

**DOI:** 10.1101/2021.11.20.469295

**Authors:** Juan Mauricio Garré, Feliksas F. Bukauskas, Michael V.L. Bennett

## Abstract

Astrocytes express surface channels involved in purinergic signaling, and among these channels, pannexin-1 (Px1) and connexin-43 (Cx43) hemichannels (HCs) mediate ATP release that acts directly, or through its derivatives, on neurons and glia via purinergic receptors. Although HCs are functional, *i*.*e*., open and close, under physiological and pathological conditions, single channel conductance of Px1 HCs is not well defined. Here, we developed a dual voltage clamp technique in HeLa cells overexpressing human Px1-YFP, and then applied this system to rodent spinal astrocytes. Single channels were recorded in cell attached patches and evoked with ramp cycles of 2 s duration and -/+ 80-100 mV amplitude through another pipette in whole cell clamp. Conductance of Px1 HC openings recorded during ramp stimuli ranged 25-110 pS. Based on their single channel conductances, Px1 HCs could be distinguished from Cx43 HCs and P2X7 receptors (P2X7Rs) in spinal astrocytes during dual voltage clamp experiments. Furthermore, we found that single channel activity of Cx43 HCs and P2X7Rs was increased, and that of Px1 HCs was decreased, in spinal astrocytes treated for 7 h with FGF-1, a growth factor implicated in neurodevelopment, repair and inflammation.

## Introduction

Pannexins (Pxs) and connexins (Cxs) are members of two different gene families that can form gap junctions (GJs) from hemichannels (HCs) in apposed membranes. In the central nervous system (CNS) these proteins are detected in neurons, glial cells, pericytes, and endothelial cells (Giaume et al., 2021; Nagy et al., 2018; Nagy and Rash, 2000; Vanlandewijck et al., 2018). Pxs and Cxs are assembled into oligomers in the ER-Golgi compartments to form HCs, which are inserted into the plasma membrane. When two compatible surface HCs dock in series, one HC from each opposed cell, they form an intercellular channel that may be incorporated into a cluster of hundreds of channels, a so-called GJ plaque. Chordate Pxs are orthologs of invertebrate innexins, the GJ forming proteins in protostomes and deuterostomes, except for echinoderms and hemichordates which have no identified GJs (Baranova et al., 2004; Bruzzone et al., 2003; Garré and Bennett, 2009; Panchin et al., 2000). There is still no evidence that vertebrate Px HCs may dock in series to form Px GJ channels *in vivo*, although they do in Xenopus oocytes (Bruzzone et al., 2003) and mammalian cell lines transfected with exogenous Px1 cDNA (Lai et al., 2007; Sahu et al., 2014; Vanden Abeele et al., 2006). Px GJ formation in mammalian tissues may be limited under physiological conditions because of glycosylation at HC docking sites (Ruan et al., 2020).

In contrast to vertebrate Pxs, Cx GJs have been shown to comprise electrical synapses between GABAergic interneurons that modulate networks of excitatory neurons in higher vertebrates such as mammals (Bennett and Zukin, 2004; Tremblay et al., 2016). In fishes, Cx GJs are found in mixed (chemical and electrical) synapses in main pyramidal neurons that control motor outputs and escape behaviors (Alcami and Pereda, 2019). A distinct function of Cx GJs in mammalian astrocytes is that GJs generate a metabolic syncytium that contributes to modulation of neural activity (Clasadonte et al., 2017; Gutnick et al., 1981; Houades et al., 2006; Rash et al., 2001; Rouach et al., 2008).

To date, several lines of evidence have shown that both Pxs and Cxs can form unopposed functional HCs in astrocytes. In general, HCs modulate a wide range of physiological and pathophysiological processes including cell death, inflammation, neurotransmission and behavior (Contreras et al., 2002; Garré et al., 2010; Iglesias et al., 2009; Orellana et al., 2011; Retamal et al., 2007; Stehberg et al., 2012). In particular, Px1 HCs have been shown to contribute to neurodegeneration and neuroinflammation (Bennett et al., 2012; Garré et al., 2016; Giaume et al., 2021; Naus and Giaume, 2016; Seo et al., 2021; Velasquez et al., 2016), and more recently, over activation of Px1 HCs has been associated with abnormal neurodevelopment and seizures (Avendano et al., 2015; Prieto-Villalobos et al., 2021; Santiago et al., 2011; Scemes et al., 2019).

Distinguishing between Px1 HC and Cx HC pathways is still a challenge, since genetic manipulations induce compensations and specific pharmacological tools are not currently available. For example, ablation of Cx43 in astrocytes is well known to affect other GJ channels (Dermietzel et al., 2000; Naus et al., 1997) and confounding behavioral phenotypes have been noted from various Px1-deficient mice (Battulin et al., 2021). Techniques that improve measurements of unitary conductance would provide a very informative tool to distinguish between Px HC and Cx HC pathways in general and in translational cellular models such as human-derived glial and neuronal cells obtained from patients with neurological and neurodevelopmental disorders.

GJ channels are nominally twice the length of one HC, and so, an intercellular channel has about twice the resistance of an unopposed HC. For example, the unitary conductance of Cx43 HCs is about twice that of the gap junction channels, *i*.*e*., ∼ 110 pS (Bukauskas et al., 2001; Bukauskas and Peracchia, 1997; Contreras et al., 2003). While early studies reported that when expressed exogenously in oocytes, human Px1 forms HCs that open with a maximum unitary conductance of ∼500 pS (Bao et al., 2004; Qiu and Dahl, 2009), others have found Px1 HCs with unitary conductances no larger than ∼ 100 pS (Chiu et al., 2017; Ma et al., 2012), and there is still no agreement on the single channel conductance of vertebrate Px1 HCs.

To clarify the unitary conductance of Px1 HCs and compare their molecular properties with those of other ion channels in astrocytes (*i*.*e*., Cx43 HCs and P2XRs), we used a dual voltage clamp technique in which current through a cell attached patch was measured during administration of a voltage applied with a whole cell voltage clamp (**Figure 1A**). After validation of the technique in HeLa cells overexpressing exogenous human Px1 tagged with yellow fluorescent protein (HeLa Px1-YFP) and rat Cx43 tagged with cyan fluorescent protein (HeLa Cx43-CFP), we recorded HCs from cultured astrocytes prepared from both embryonic and neonatal rodent spinal cords, which are known to express endogenous Px1 and Cx43 (Garré et al., 2010). Px1 HCs were activated by V_m_ with voltage ramp cycles of 2 s duration and -/+ 100 mV amplitude in HeLa Px1-YFP cells. We found that Px1 HCs open with unitary conductances ranging from 25 to 110 pS. Single channels with unitary conductances similar to those obtained in HeLa Px1-YFP cells were recorded in rodent spinal astrocytes, and differed significantly from those of Cx43 HCs (∼160-250 pS) and P2X7Rs (∼17 pS). Furthermore, in astrocyte cultures treated with fibroblast growth factor 1 (FGF-1) for 7 h, the percentage of ramps with active Px1 HCs was decreased, whereas that of P2X7Rs and Cx43 HCs was increased.

**Figure 1.**
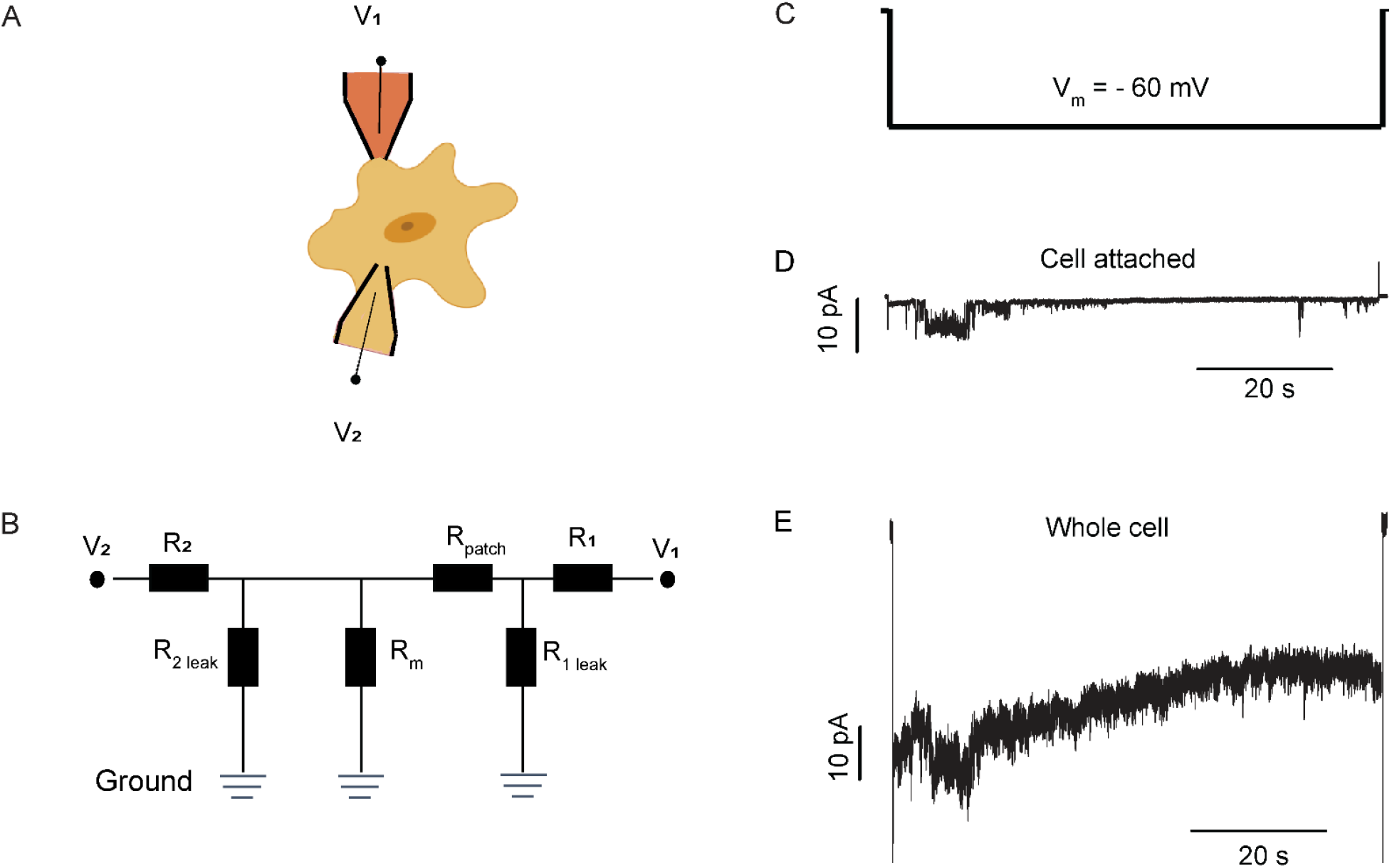
Validation of dual voltage clamp technique in HeLa Px-1 YFP cells. (**A, B**) Diagram (A) and circuit (B) of dual patch clamp with one pipette in cell attached mode (V_1_) and one pipette in whole cell mode (V_2_). V_1_, recorded by cell attached pipette. R_1_, V_1_ electrode resistance. R_patch_, patch resistance. R_1 leak_, leak around patch to bath. V_2_, recorded by whole cell pipette. R_2_, V_2_ electrode resistance. R_2 leak_,, leak around V_2_ electrode where it enters the cell. R_m_, whole cell membrane resistance. (**C**) Voltage pulse applied through the whole cell pipette. (**D**) Single channel currents recorded with the cell attached pipette using 50 mV/pA gain. (**E**) Recording with the whole cell pipette at the same gain as that used in (**D**). Currents were recorded simultaneously at the same gain in the cell attached patch (**D**) and whole cell pipettes (**E**); V_m_ = (V_2_ – V_1_) was held at –60 mV. To present inward currents in the same direction in both whole cell and cell attached patch recordings, actual currents recorded through the V_1_ electrode were multiplied by -1.

## Results

### Unitary events recorded from Px1-YFP expressing HeLa cells

HeLa cells overexpressing human Px1-YFP (HeLa Px1-YFP) or rat Cx43-CFP (HeLa Cx43-CFP) (**Figure S1**) offer simple models to record unitary events of Px1 HCs and Cx43 HCs, respectively, with less interference from activity of other ion channels. Using HeLa Px1-YFP cells, we improved signal to noise ratio and calculation of unitary conductances in single channel recordings by using a dual patch clamp technique. In the dual patch recordings, one pipette was in cell attached mode to record activity of single channels in the limited area of the patch, and the other was in whole cell mode to clamp V_m_. The whole cell pipette was used for both stimulating and recording whole cell activity, including that of channels in the cell attached patch (see diagram in **Figure 1A, B**). The signal to noise ratio in single channel recordings was greater in the cell attached patches than in the whole cell recordings (**Figure 1C-E**).

HC openings as in **Fig. 2** were evoked with ramps of 1.7 s duration and -/+ 100 mV amplitude. Ramps started with a fast hyperpolarization from 0 to -100 mV, where V_m_ (V_2_ - V_1_) remained at – 100 mV for 0.1 s. This was followed by a linear rise in V_m_ from - to + 100 mV over 1.5 s, V_m_ remained stationary again for another 0.1 s. Ramp cycle ended with a fast return to 0 mV, where V_m_ remained for 0.3 s (total cycle time: 2 s, **Figure 2A**). Channel openings were detected in cell attached patches during ramp stimuli, and were simultaneously recorded, at lower resolution, through the whole cell pipette (**Figure 2B-D**). Ramp-evoked HC openings showed a broad range of unitary conductances (25-110 pS). The mean unitary conductance from these distributions recorded during ramps was 50.8 ± 4.2 pS and 48.3 ± 3.8 pS, at negative and positive V_m_, respectively (*P* = 0.657) (**Figure 2E**).

**Figure 2.**
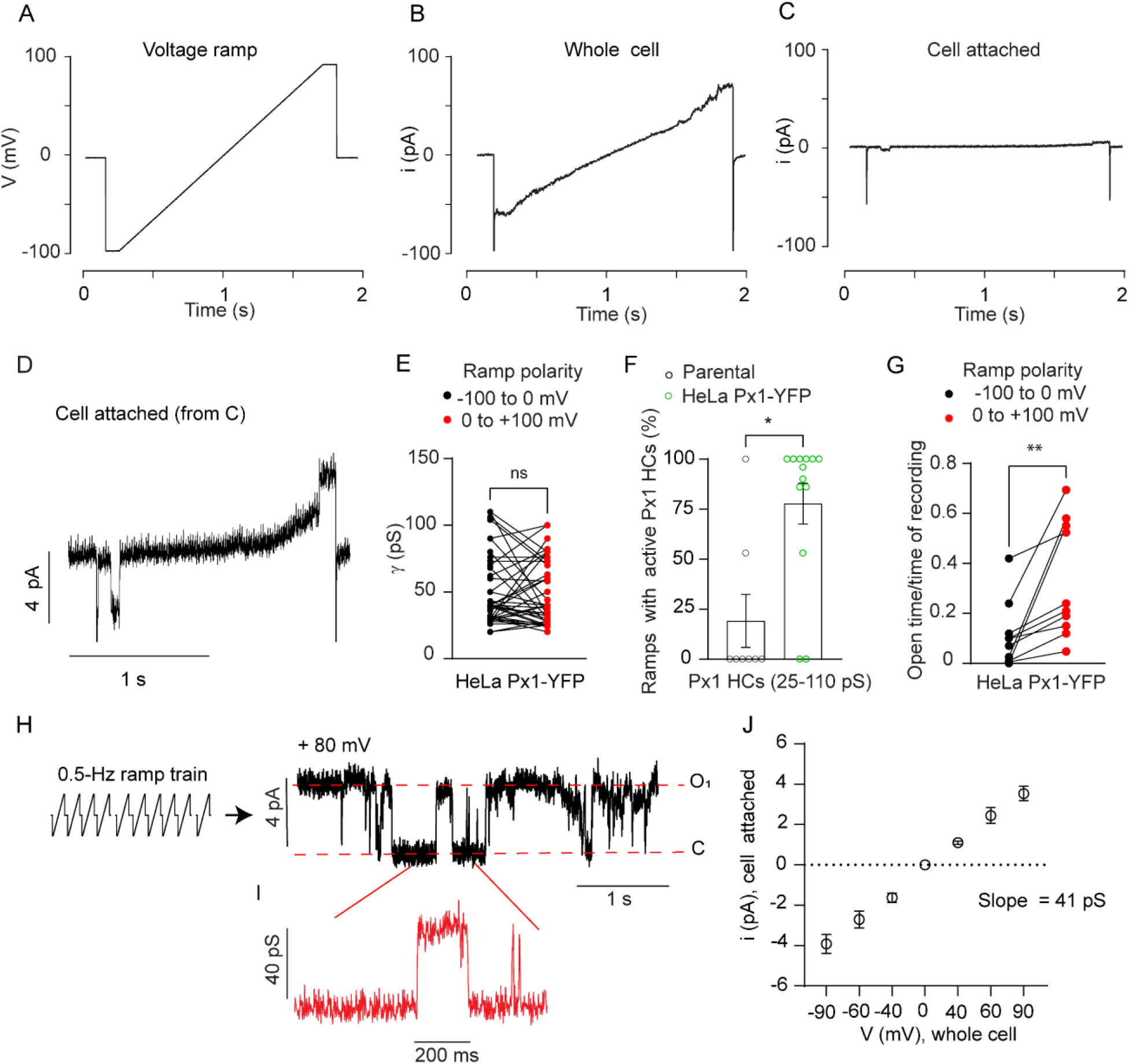
Single channels recorded using dual voltage clamp in HeLa Px1-YFP cells. (**A-C**) Ramps of 1.7 s duration and −/+ 100 mV amplitude applied to HeLa Px1-YFP cells (**A**) evoked (hemi)channel openings recorded in both whole cell (**B**) and cell attached patch recordings (**C**). Currents were recorded simultaneously in the cell attached patch (**C**) and whole cell pipettes (**B**) at a gain of 50 mV/pA. (**D**) Same cell attached patch recording shown in **C**, but at higher current amplification. (**E**) Quantification of single channel conductances. Unitary conductances were calculated by identifying channel opening and closing during the linearly changing phase of voltage ramps from experiments as shown in **D**. Single channel conductances were also confirmed from the I-V curves from the same experiments (*n* = 40 ramps, 5 ramps/patch, from 8 different patches and cells, mean = 50.8 ± 4.2 pS, V_m_ from -100 to 0 mV, mean = 48.3 ± 3.8 pS, V_m_ 0 to 100 mV). (**F**) Percentage of ramps with active channels in cell attached patches (*i*.*e*., fraction of ramps with active channels, 30 consecutive ramps as in panel A for each cell) in HeLa parental or Px1-YFP cells (*n* = 8, 13 patches, one patch per cell). (**G**) Total open time divided by total period of recording during ramp stimulation (*n* = 10 patches, one patch per cell). (**H**) After evoking Px1 HC openings with a 0.5-Hz train of voltage ramps, V_m_ was clamped at +80 mV through a whole cell pipette and single channel activity was recorded in the cell attached pipette. C, closed state, O_1_ open state. (**I**) Unitary conductance calculated from experiments like those in **H**. (**J**) Single channel I-V curves (*n* = 12 voltage pulses, 2 pulses per patch for each voltage, from 6 different HeLa Px1-YFP cells). Both pipettes contained internal recording solution because the cell attached patch was opened at the end of the experiment to calculate the voltage drop in the V_2_ electrode (see Methods). MgATP was not added to the cell attached pipette solution to minimize activation of purinergic receptors. Data are presented as dot plots in (**E, G**) and mean ± SEM in (**F**). \**P* < 0.05, ** *P* < 0.01, Wilcoxon matched-pairs signed rank test was used in **E, G**, and Mann Whitney test was used in **F**.

Voltage ramps applied through the whole cell pipette produced channel openings in cell attached patches of HeLa Px1-YFP cells at higher prevalence (∼78%) than in patches of parental cells (∼19%), indicating a higher number of active HCs in the cell surface of HeLa Px1-YFP, as compared to parental cells (**Figure 2F**) (*P* < 0.05). The mean open time of single Px1 HC openings during 15 consecutive ramps (30 s) at positive V_m_ was greater than at negative V_m_ (**Figure 2G**, *P* < 0.05).

Single channel conductance of unitary events was determined in HeLa Px1-YFP cells when V_m_ was held through the whole cell pipette at, for example, + 80 mV, and after activation of HCs with a train of at least 10 consecutive voltage ramps of 2 s cycle time and -/+ 100 mV amplitude (**Figure 2H**,**I**). Active channels in HeLa Px1-YFP cells showed a near linear single channel I-V relationship in which the mean slope was 41 pS (**Figure 1J**).

Together these data demonstrate that a dual voltage clamp technique can be used to characterize ion channels (*e*.*g*., Px1 HCs) in HeLa cells with improved recording of unitary events.

### Pharmacological block of Px1 HCs in HeLa Px1-YFP cells

To confirm that Px1 HCs mediate unitary currents recorded from cell attached patches in HeLa Px1-YFP cells, we inhibited channel activity pharmacologically. Agents in the cell attached pipette rapidly reach the target site in the cell membrane under the pipette. We added hemichannel blockers to the cell attached pipette solution and measured the channel activity and conductance in the patch 1 min after gigaseal formation. Channels were recorded as in **Fig. 2D**. The electrical conductance in the cell attached patch was calculated by measuring the slope of the I-V curve, where I in this case, was the current flowing through the cell attached pipette, and V_m_ was the voltage clamped through the whole cell pipette (V_2_ - V_1_). The conductances of the patches of HeLa Px1-YFP cells were significantly greater than those of cell attached patches of HeLa Px1-YFP cells to which CBX (0.05 mM) or probenecid (1 mM), two Px1 HC blockers, were applied (**Figure 3A-C**). CBX is a well-known gap junction and Cx HC blocker that also affects Px HCs (Garré et al., 2010), whereas probenecid blocks Px HCs with little effect on Cx HCs (Silverman et al., 2008). CBX and probenecid blocked more completely during the negative phase of the ramps than during the positive phase (**Figure 3B, C**) (negative phase, *P* < 0.001 probenecid and CBX vs. control; positive phase, *P* < 0.001 and *P* < 0.01, probenecid and CBX vs. control, respectively). Presumably, additional kinds of channels contributed to the positive phase. Furthermore, the maximum number of channel openings was reduced in patches containing CBX and Probenecid, as compared to untreated cells (**Figure 3D**) (*P* < 0.05). Together, these data confirm that the major single channel activity recorded in the cell attached patches in HeLa Px1-YFP cells was mediated by Px1 HCs.

**Figure 3.**
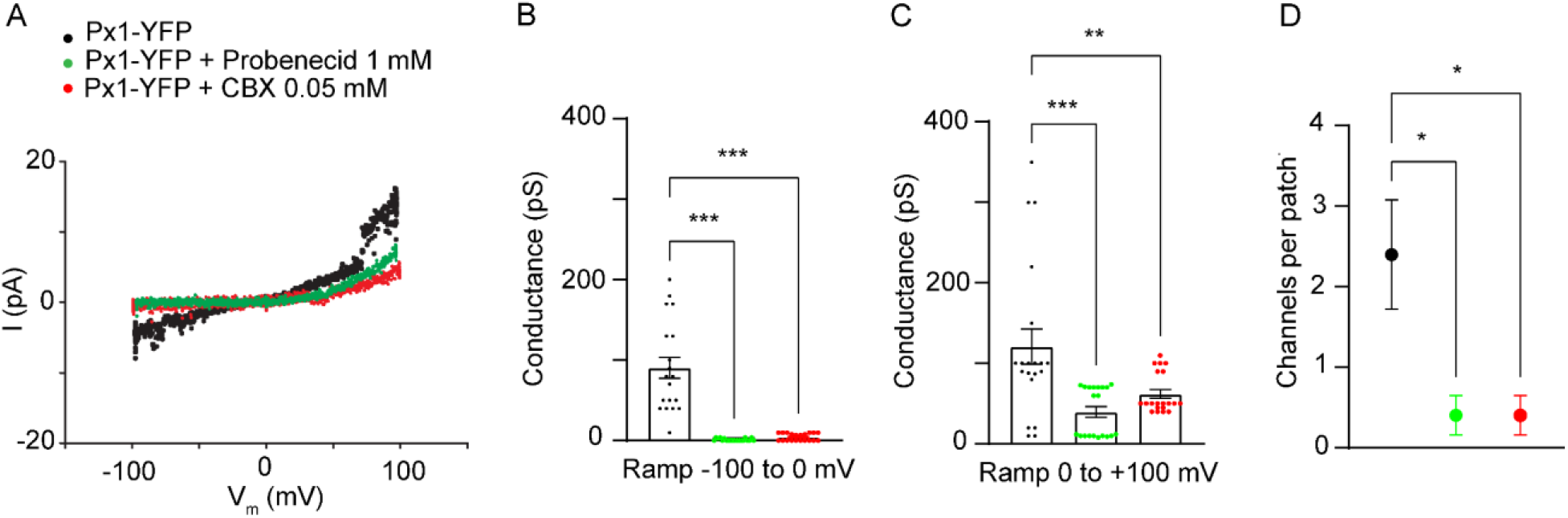
Pharmacological block of Px1 HCs in cell attached patches. (**A**) Px1 HCs are sensitive to carbenoxolone (CBX) and probenecid. CBX also blocks Cx HCs. Probenecid is a Px1 HC blocker with no effect on Cx HCs. In these I-V curves like those in **Fig. 2**, I is the current in the cell attached pipette, and V_m_ is the cell membrane voltage which was clamped through the whole cell pipette (V_m_ = V_2_-V_1_). (B) Conductance (pS) in the patch during the more negative phase of the voltage ramp (−50 to -100 mV) in HeLa Px1-YFP control and treated with probenecid (1 mM) or CBX (50 μM) (*n* = 20 ramps, from 4 different patches and cells, probenecid and CBX vs. control). (C) Conductance in the patch during the positive phase of the voltage ramp (+50 to 100 mV) (*n* = 20 ramps, from 4 different patches and cells, probenecid and CBX vs. control, respectively). (D) Maximum number of channel openings observed per patch (*n* = 5 patches, 10 ramps per patch, one patch per cell). Drugs were added to the cell attached pipette solution. Data are presented as dot plots and the mean ± SEM. \**P* < 0.05, \*\*\**P* < 0.001, one-way ANOVA followed by Sidak’s multiple comparisons test was used in **B, C, D**.

### Unitary currents in whole cell recordings from HeLa Px1-YFP cells

Mechanical stress on the cell surface might increase voltage sensitivity of channels in the patch, and cell dialysis caused by the whole cell pipette could affect P_o_. In separate experiments, we monitored single channel activity in whole cell recordings, but not using a second, cell attached pipette. V_m_ was clamped at +/-70 and +/-40 mV for ∼10 s, somewhat longer steps than the ramp stimuli that we used in **Fig. 2** (**Figure S2A-F**). Single channels opened with mean unitary conductance of 32.4 ± 9.8 pS, 35.9 ± 8.2 pS, 37.9 ± 9.3 pS, and 31.1 ± 9.1 pS, at 70, 40, -40, -70 mV (*P* > 0.05). These events had linear I-V relationships with slopes of 31 pS and 44 pS at negative (−50-100 mV) and positive (50-100 mV) V_m_, respectively (**Figure S2F**), values within the range of conductances recorded during ramp stimuli (25-110 pS) and voltage steps using dual voltage clamp (∼40 pS, see **Figure 2I**).

These data indicate that single channels recorded from HeLa Px1-YFP cells using whole cell recordings show single channel conductances similar to those of Px1 HC openings recorded using a dual voltage clamp technique. Furthermore, in contrast to a previous report using oocytes (Bao et al., 2004), we found no evidence of channels of larger electrical conductance, *e*.*g*., ∼ 500 pS, in HeLa Px1-YFP cells in either dual voltage clamp or whole cell recordings.

### Unitary events recorded from Cx43-CFP expressing HeLa cells

In HeLa cells transfected with WT Cx43 or Cx43-EGFP, HCs open at low probability and with a maximum single channel conductance of ∼220 pS (Contreras et al., 2003). We failed to see the predicted single channel activity of Cx43 HCs (∼220 pS) in Cx43-CFP transfected HeLa cells with a dual voltage clamp and ramp stimulation as used for Px1 HCs (2 s ramp cycles of -/+ 100 mV amplitude, **Figure 4A, 4B, and 4F**). However, short ramps evoked discrete channel openings ranging from 6 to 80 pS (19.3 ± 6.8 pS), likely mediated by endogenous Cx45 HCs or other channels (Choi et al., 2020). Cx43 HC-like unitary currents were observed during 8 s ramp cycles of the same amplitude, mostly at positive voltages (**Figure 4C-F**), indicating that Cx43 HC P_o_ at negative V_m_ is very low. In HeLa Cx43-CFP cells, the mean unitary conductance of channels in the range of Cx43 HCs (*i*.*e*., 160-250 pS) was 180.9 ± 8.1 pS (**Figure 4F**). During 8 s ramp stimulation, there were also channel openings of lower unitary conductance ranging from 80 to 150 pS (109.2 ± 6.9 pS), likely representing Cx43 HC substates or other channels.

**Figure 4.**
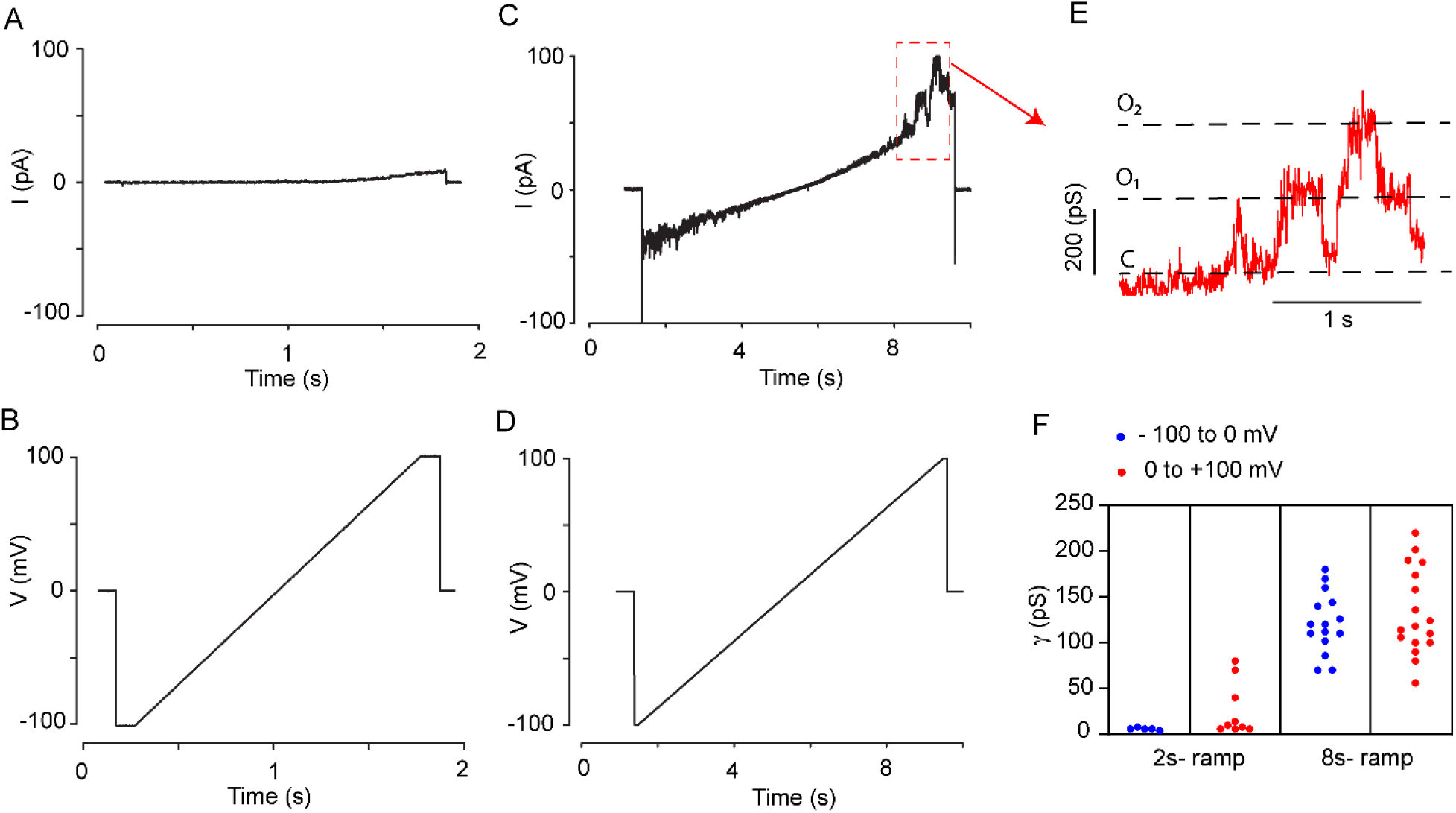
Single channels recorded using dual voltage clamp in HeLa Cx43-CFP cells. (**A, B**) Ramps of 1.7 s duration (2s-ramp) and −/+ 100 mV amplitude applied to HeLa Cx43-CFP cells did not evoke (hemi)channel opening in the cell attached pipette (A). Voltage ramps (B) were applied through the whole cell pipette, while V_2_ in the cell attached pipette was held at 0 mV. (**C, D**) Ramp of 7.7 s duration (8s-ramp) and −/+ 100 mV amplitude (**D**) evoked Cx43 hemichannel opening in the patch (**C**). (**E**) Calculated single channel conductances from channel transitions observed at positive V_m_ in traces like those in **C**. C, closed state. O_1_ and O_2_ one, two channels open, respectively (**F**) Single channel conductances from experiments using short (1.7 s) and long (7.7 s) duration ramps (2s-ramp cycle, *n* = 6-9 ramps, 3 patches, one patch per cell; 8s-ramp cycle, *n* = 14-16 ramps, 3 patches, one patch per cell).

### Single channel properties of Px1 HCs and Cx43 HCs in rodent spinal astrocytes

In contrast to HeLa cells (**Figure S2**), our whole cell recordings in astrocytes did not show a signal to noise ratio adequate to discriminate small transitions ranging between ∼20 pS and ∼40 pS, which would include unitary conductances for both P2X7Rs and Px1 HCs, respectively. Having used a dual voltage clamp technique in HeLa cells to record HCs, we asked whether the same technique would allow activity of these channels to be distinguished from that of Cx43 HCs and P2XRs in rat and mouse astrocytes.

We applied ramp stimulation (2 s ramp cycle, stimulation 1.7s, -/+ 80 mV) through a whole cell clamp and recorded channel activity in cell attached patches in spinal astrocyte cultures (**Figure 5A**). We first assigned channel activity to Px1 HCs, Cx43 HCs or P2X7Rs based on the expected unitary conductances reported in the literature or from our recordings in HeLa cells and astrocytes (*i*.*e*., Px1 HCs = 25-110 pS, Cx43 HCs = 160-250 pS, P2X7Rs = 10-22 pS). We confirmed these data by pharmacological treatment and genetic depletion of Px1 and Cx43.

**Figure 5.**
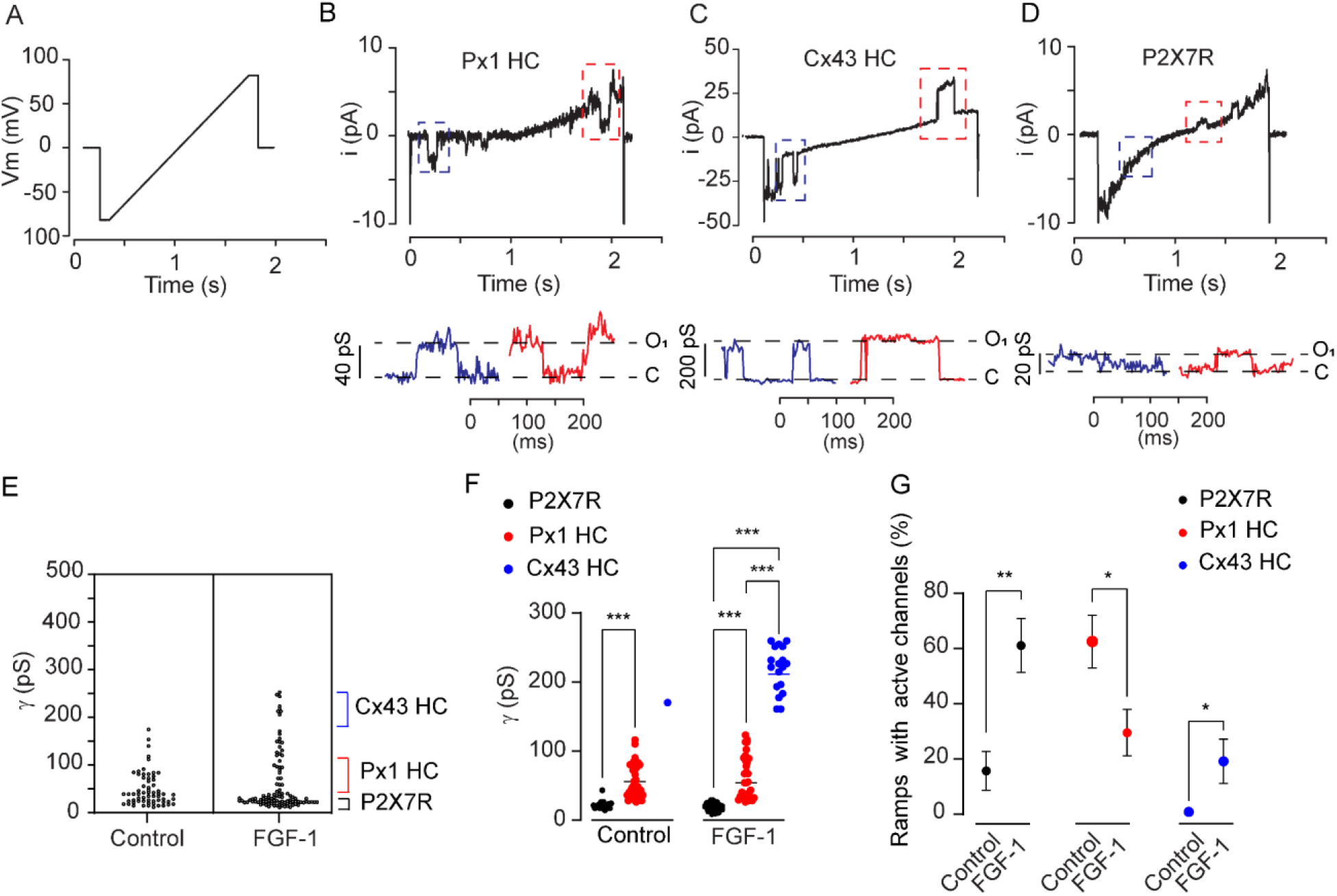
Single channel transitions in spinal astrocytes cultured from neonatal rat spinal cords. (**A**) Channel transitions were recorded in cell attached patches during ramps of 1.7 s duration and -/+ 80 mV amplitude applied through a whole cell pipette. (**B-D**) Representative single channel recordings of Px1 HCs (**B**), Cx43 HCs (**C**), and P2X7Rs (**D**) in rat spinal astrocytes after 7 h FGF-1 treatment. (**E**) Dot plot distribution of single channel conductances in astrocytes, control and treated 7 h with FGF-1 (*n* = 60 ramps, 6 patches, one patch per cell). (**F**) Distributions of active P2X7Rs, Px1 HCs and Cx43 HCs in astrocytes maintained in control solution and after treatment for 7 h with FGF-1. Events were grouped in 3 statistically different categories: unitary single channel conductances of 10-22 pS (P2XRs including P2X7Rs), 25-110 pS (Px1 HCs), 160-250 pS (Cx43 HCs) (*n* = 18-36, *n* = 62, 36, *n* = 36, 18, *n* = 62, 18, patches, one patch per cell). Unitary events < 10 pS and ranging from 120 to 150 pS were undetermined. (**G**) Percentage of ramps with active P2X7Rs, Px1 HCs, and Cx43 HCs in control and 7 h FGF-1-treated astrocytes. After 7 h treatment with FGF-1, the percentage of ramps with active P2X7Rs and Cx43 HCs increased, and with Px1 HCs decreased (*n* = 14, 18, *n* = 11, 20, *n* = 12, 12, patches, one patch per cell). Data are presented as dot plots in **E, F**. The mean ± SEM is included in **F, G**. \**P* < 0.05, \*\**P* < 0.01, \*\*\**P* < 0.001, one-way ANOVA followed by Sidak’s multiple comparisons test was used in **F** for control and FGF-1, Mann Whitney test was used in **G**.

Representative traces for Px1 HCs, Cx43 HCs, and P2X7Rs recorded in spinal astrocytes treated 7 h with FGF-1 are shown in **Figure 5B-D**. The amplitude and frequency of unitary events were changed after treating astrocytes for 7 h with FGF-1, as compared to astrocytes maintained in control medium (**Figure 5E**). This is consistent with a previous study showing that FGF-1 increases ATP release and opens hemichannels and P2X7Rs in spinal astrocytes (Garré et al., 2010). The mean unitary conductances of the single channel events assigned to Px1 HCs, Cx43 HCs and P2X7Rs were significantly different from each other in both control and FGF-1-treated astrocytes (**Figure 5F**) (*P* < 0.001). The percentage of ramps with active Cx43 HCs was increased in astrocytes that had been in culture medium with FGF-1 for 7 h, as compared to astrocytes maintained for 7 h in control medium alone (*P* < 0.05). Although a fraction of ramps showed Px1 HC openings after 7 h FGF-1 treatment (∼ 30%), the percentage of ramps with active Px1 HCs was reduced as compared to control astrocytes (*P* < 0.05). Both Px1 HC and Cx43 HC openings mediate ATP release, which in turn activates P2X7Rs. Accordingly, the percentage of ramps with active P2X7Rs was increased after 7 h FGF-1 treatment, as compared to control astrocytes (**Figure 5G**) (P < 0.01).

To confirm the identity of HCs, we conducted dual patch clamp experiments in Px1-deficient and Cx43^-/-^ astrocytes. The percentage of ramps with active 25-110 pS channels was reduced in spinal astrocytes transfected with stealth siRNA against Px1 (siRNA Px1), as compared to astrocytes transfected with a non-targeting siRNA (siRNA(-)), indicating that these unitary events were mostly mediated by Px1 HCs (**Figure 6A-C**). Of note, siRNA Px1 treatment did not prevent openings of < 25 pS channels, indicating this activity is not mediated by Px1 HCs. Furthermore, we observed possible compensatory effects on the single channel activity of other channels including P2X7Rs (10-22 pS) and Cx43 HCs in Px1-deficient spinal astrocytes (**Figure 6A**).

**Figure 6.**
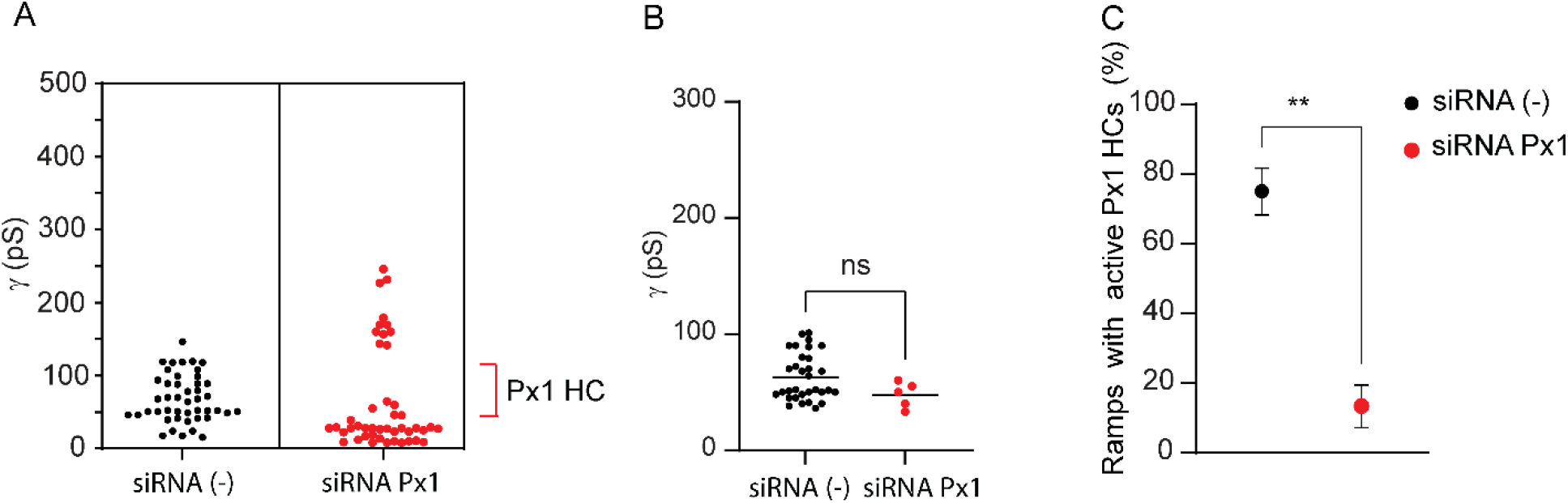
Px1 knockdown in rat spinal astrocytes reduces the number of channel transitions in the range of Px1 HCs (25-110 pS). (**A**) Single channels were recorded in the cell attached pipette whereas V_m_ was controlled through a whole cell clamp after 24 h transfection of rat spinal astrocytes with siRNA (−) and siRNA Px1. Single channel transitions were evoked by voltage ramps of 1.7 s and -/+ 80 mV. (**B**) Mean unitary conductance of Px1 HCs was not altered after transfection with siRNA (−) and siRNA Px1 (*n* = 32, *n* = 5, ramps with 25-110 pS channels, *P* > 0.05). (**C**) Percentage of ramps with active Px1 HCs (25-110 pS) was reduced after 24 h transfection with siRNA Px1 (*n* = 45 ramps, 5-10 ramps/patch, 6 patches, one patch per cell). Data are presented as dot plots in **A**, B. Mean ± SEM is included in **B, C**. \*\**P* < 0.01, ns, not significant, Mann Whitney test was used in **B, C**.

We also recorded Px1 HC openings (∼ 40 pS) at constant V_m_ in astrocytes prepared from Cx43^-/-^ mouse embryos. The channel activity of 40 pS was unlikely to be mediated by Cx30 HCs, which are expected to form HCs of larger unitary conductance (∼300 pS) and are not detected in the CNS until P10 (Ribot et al., 2021). This ∼ 40 pS unitary activity was reduced after bath application of CBX (200 μM), confirming that these channel openings were primarily mediated by Px1 HCs (**Figure 7**). The distribution of channel conductances in mouse astrocytes maintained in control medium was similar to that in rat astrocytes (**Figure 7** and **Figure S3**). As expected, Cx43 HC openings (160-250 pS) were not observed in spinal astrocytes prepared from Cx43 KO mouse embryos (**Figure 7B** and **Figure S3**). Overall, Cx43 HCs openings were rare in control rat and mouse astrocytes, indicating that in the cell attached patch, Cx43 HC P_o_ was very low.

**Figure 7.**
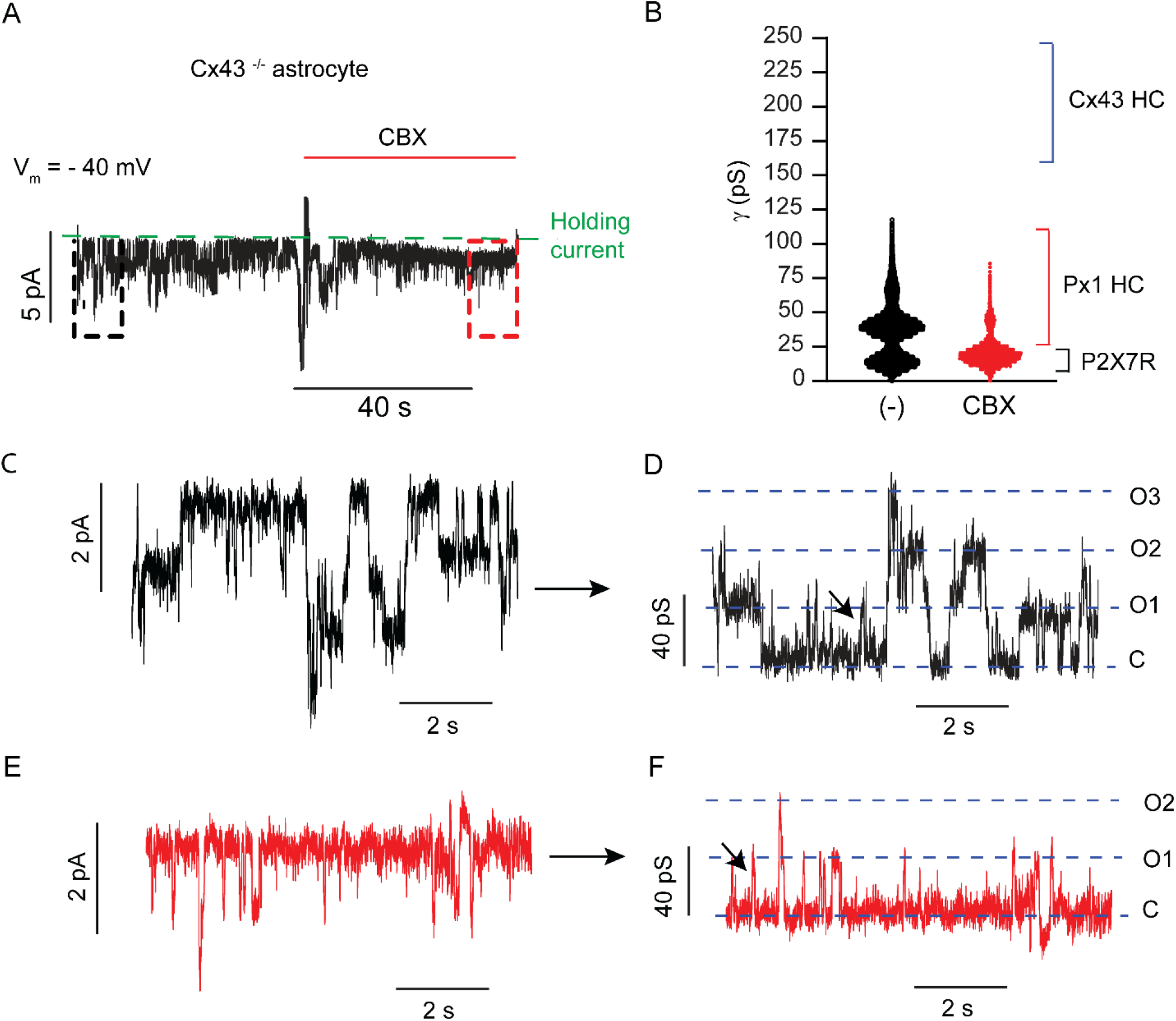
Px1 HCs are functional in spinal astrocytes prepared from Cx43 KO mouse embryos. (**A**) Single channel currents before and during bath application of carbenoxolone (CBX) were recorded in the cell attached pipette. V_m_ was clamped with a second whole cell pipette at - 40 mV. After putative Px1 HC activity was recorded (∼ 40 pS), the Px1 HC and Cx43 HC blocker CBX (200 μM) was applied to the bath to confirm channel identity. Holding current is indicated with a green dotted line. (**B**) Single channel distribution 40 s before (black) and 40 s after (red) CBX application. CBX reduced single channel transitions in the range of Px1 HC (25-110 pS). Single channel transitions in the range of P2X7Rs were not affected by CBX. Single channel transitions in the range of Cx43 HCs were not detected in Cx43^-/-^ mouse spinal astrocytes. (**C-F**) Single channel currents (**C, D**) and unitary conductances (**E, F**) from the traces in **A** before (**C, E**) and after (**D, F**) application of CBX. C, closed state. O1, O2, O3, one, two, and three Px1 (hemi)channels opened, respectively. Single channel transitions in the range of P2X7Rs (10-22 pS) are indicated with black arrows in (D, F).

In rat and mouse spinal astrocytes, we found another type of high unitary conductance channel; these channels opened with unitary conductances ranging 120-150 pS in astrocytes in both control and after 7 h FGF-1 (**Figure 5E** and **Figure S3)**. These transitions were unlikely to mediated by Px1 HCs and Cx43 HCs, because they were not observed in HeLa Px1-YFP cells (**Fig. 2E**), but were often detected in patches from Px1-deficient and Cx43^-/-^ astrocytes (**Figure 6A** and **Figure S3**). Further studies will be required to determine the molecular basis of these channel openings.

### Single channel properties of P2X7Rs in rodent spinal astrocytes

Unitary transitions ranging from 10 to 22 pS resemble the single channel responses of P2XRs including P2X7Rs (Benham and Tsien, 1987; Riedel et al., 2007). The mean unitary conductances of P2X7Rs calculated from dual voltage clamp experiments (as those shown in **Figure 5D**) in rat spinal astrocytes in control and after 7h FGF-1 treatment were 16.9 ± 3.5 pS and 17.3 ± 4.8 pS, respectively. P2X7R single channel events show a non-linear current-time dependence. The non-linearity in the base line current during the ramp was more evident as the polarity increased (−50 to -100 mV and + 50 to +100 mV, **Figure 5D**).

We also used cell attached patch single clamp (*i*.*e*., without controlling V_m_ through a second pipette) to confirm that ATP induces unitary events in the range of conductances for P2X7Rs in spinal astrocytes. Small unitary events were more frequent in patches when 50 μM or 500 μM ATP was added in the recording solution, as compared to control patches without ATP (**Figure S4A-C**). Unitary events were ∼2.5 pA at V_p_ = + 90 mV, corresponding to unitary conductances of ∼ 22 pS (the mean resting potential of cultured astrocytes recorded in separate experiments using current clamp mode in whole cell recording was - 21.14 ± 2.58 mV). The incidence of channel openings observed was higher in the presence of ATP than in its absence (ATP 500 μM: ∼4 channels/patch; ATP 50 μM: ∼2 channels/patch, no ATP: ∼1 channel/patch).

To confirm activation of P2X7Rs by ATP, we also used cell attached single clamp in spinal astrocytes in control and after 7 h FGF-1 treatment but adding ATP to the cell attached patch recording solution (**Figure S4D**). In control astrocytes, the percentage of trails (voltages pulses, + 90 mV relative V_m_, ∼10 s duration) with active P2X channels increased in experiments in which 50 μM (41.54 ± 6.59 %) and 500 μM ATP (33.85 ± 7.30 %) was added to the cell attached pipette (50 μM, *P* < 0.05, 500 μM, *P* = 0.05), as compared to results in the absence of ATP (9.23 ± 4.31%). Similarly, in astrocytes treated with FGF-1 for 7 h, the percentage of trials with active P2X7Rs increased when ATP (50 μM) was present in the recording solution (33.84 ± 6.94 %), as compared to control astrocytes in which ATP was not added to the cell attached pipette (*P* = 0.05) (**Figure S4E**).

Furthermore, when a P2X7R antagonist, Brilliant Blue G (BBG, 10 μM) was added together with ATP in the cell attached pipette, events of < 25 pS were absent in both control astrocytes and those treated with FGF-1 for 7 h. Moreover, at this high concentration of BBG, Px1 HCs and Cx43 HCs were functional in both control and FGF-1 treated astrocytes (**Figure S4I**). These data indicate that unitary currents (∼22 pS) induced by high ATP concentrations (50, 500 μM) are primarily mediated by P2X7Rs, although other channels such as P2X2R and P2X4R may contribute.

Together, these data show that Px1 HCs, Cx43 HCs and P2X7Rs are functional in astrocytes prepared from embryonic and neonatal rodent spinal cords. These channels can be distinguished by their single channel conductances, gating, and pharmacological properties, and their single channel activities are modulated by FGF.

## Discussion

We developed a dual voltage clamp technique to reveal the properties of gap junction hemichannels in cultured primary spinal astrocytes. We focused on describing the single channel properties of Px1 HCs at first, and then, compared it with Cx43 HCs and P2XRs expressed by spinal astrocytes. Px1 HCs were functional and could be differentiated at the single channel level from Cx43 HCs and P2X7Rs using this dual voltage clamp technique. Px1 HCs openings showed single channel conductances ranging over 25-110 pS. Single channel activity of HCs and P2X7Rs in spinal astrocytes was modulated by FGF-1, suggesting that these surface channels may be involved in neurodevelopment, repair and spinal cord inflammation.

There is no agreement as yet on the single channel conductance of Px1 HCs. Previous reports stated that the maximum unitary conductance of Px1 HCs in oocytes was ∼500 pS. These data led to the idea that vertebrate Px1 HCs may form “pores” of large electrical conductance in the membrane. These pores should be big enough to accommodate large intracellular molecules and dyes, such as ATP, IL1-β and YO-PRO (Bao et al., 2004; Pelegrin and Surprenant, 2006), and if activated in neurons, the opening of these large channels might drive circuit dysfunction (Thompson et al., 2008). A later study suggested Px1 HCs form anion selective channels of smaller unitary conductance (∼ 70 pS) in mammalian cell lines, as evaluated in outside out patches (Ma et al., 2012). To explain this disparity, it has been also suggested that Px1 may have two different open channel configurations depending on the concentration of extracellular potassium, [K]_o_. The two different conductances of the Px1 HCs may be associated with different channel permeabilities to ATP. A relatively small channel of 50 pS would likely be ATP-impermeable, whereas, a larger channel of ∼500 pS (activated by high [K]_o_) would permit ATP efflux (Wang et al., 2014). In disagreement with these ideas, a recent study has shown that the single channel conductance of Px1 HCs is far below ∼500 pS, and channel openings of lower unitary conductances may potentially mediate ATP permeation through the channel (Chiu et al., 2017). This study, which was also conducted in outside out patches, describe a caspase-dependent mechanism of Px1 HC activation, with channel openings of 12 pS and 96 pS, at negative and positive V_m_, respectively.

To record activity of cell surface channels in intact cells with high resolution and precision we conducted dual voltage clamp experiments. In some experiments, as controls, we utilized other conventional patch clamp approaches such as single clamp cell attached patch and whole cell recordings. Unitary conductances of human and rodent Px1 HCs during ramp stimuli ranged from 25 to 110 pS. We confirmed that these unitary currents were mediated by Px1 HCs in HeLa Px1-YFP cells by pharmacology, *i*.*e*., they were blocked by probenecid or carbenoxolone. Using whole cell recordings in HeLa Px1-YFP cells, HC openings at constant negative V_m_ were of slightly lower conductance than in dual voltage clamp experiments, *i*.*e*., ∼ 30 pS, but within the range of conductances recorded during ramps. In dual voltage clamp recordings, the single Px1 (hemi)channel I-V curves show no signs of outwardly rectification, which is in contrast to the strong rectification observed in outside out patches after caspase activation in (Chiu et al., 2017). Futures studies using dual voltage clamp during caspase-induced Px1 HC activation would help to clarify this discrepancy.

Using dual voltage clamp, we identified Px1 HC openings in spinal astrocytes. Channel identity was confirmed in Px1-deficient spinal astrocytes. Importantly, we found no evidence of pores of higher electrical conductance, such as ∼ 500 pS, in either HeLa cells or rodent spinal astrocytes using dual voltage clamp and single voltage clamp whole cell and cell attached patch recordings.

There is less debate about the single channel properties of Cx43 HCs. A previous study has shown that Cx43 HCs open and close with transitions of ∼ 220 pS (Contreras et al., 2003). Accordingly, in our experiments using dual voltage clamp in HeLa Cx43-CFP and spinal astrocytes, Cx43 HCs open with distributions ranging from 160 to 250 pS These unitary currents were absent in Cx43-null spinal astrocytes.

HCs mediate ATP release from astrocytes (Abudara et al., 2018; Kang et al., 2008; Orellana, 2016), which in turn activates various ATP-gated ion channels (P2XRs) and G-protein coupled receptors (P2YRs) expressed by neurons and various glial cell types (Khakh and North, 2012). Interestingly, our dual voltage clamp technique also allowed us to record ion channels of smaller conductances (∼ 17 pS) in cultured spinal astrocytes. We confirmed that these small unitary currents were primarily mediated by P2X7Rs using cell attached patch clamp and pharmacology, *i*.*e*., activating single channels in the patch with ATP and blocking them with BBG, a P2X7R antagonist.

FGF signaling has multiple roles in normal brain and spinal cord development (Diez Del Corral and Morales, 2017; Ford-Perriss et al., 2001). Accordingly, dysregulation in various FGF systems has been associated with neurodevelopmental and psychiatric disorders (Turner et al., 2016). In the adult CNS, FGF-1 suppresses astrocyte activation after brain injury (Kang et al., 2014). Although FGF1 and FGF2 promote axonal growth and functional recovery in spinal cord injury (SCI) models, in Amyotrophic Lateral Sclerosis, FGFs favor motor neuron death and accelerate disease onset, suggesting a bimodal role of FGF signaling in inflammatory spinal cord diseases (Cassina et al., 2005; Klimaschewski and Claus, 2021). A mechanism underlying the pro-inflammatory effects of FGF-1 consists in promoting ATP release from astrocytes through Px1 HCs and Cx43 HCs (Garré et al., 2010; Garré et al., 2016). Massive release of ATP, as occurs at early stages after injury (Wang et al., 2004) may activate P2X7Rs not only in glia, but also in infiltrating inflammatory cells (Di Virgilio et al., 2017). Our data provide molecular evidence that FGF-1 increases the single channel activity of P2X7Rs in astrocytes, likely, as a consequence of the opening of Px1 HCs and Cx43 HCs, which mediate ATP release (Garré et al., 2010; Kang et al., 2008). The activation of, Px1 HCs, Cx43 HCs and P2X7Rs has been shown to self-regenerate ATP release from spinal astrocytes (Garré et al., 2016). Furthermore, ATP release may activate P2X7Rs in other types of cells than astrocytes, mediating neuronal dysfunction in neurodegeneration (Wang et al., 2004) and neurodevelopmental disorders (Garré et al., 2020).

Notably, we found that single channel activity of Px1 HCs was decreased by 7 h FGF-1 treatment in spinal astrocytes. However, a fraction of Px1 HCs remained active after this treatment (∼ 30 %), which may explain the Px1-dependent mechanism of ATP release and dye uptake observed after 7 h FGF-1 in a previous study (Garré et al., 2010). Furthermore, ATP release from astrocytes can be achieved by opening of Cx43 HCs. Indeed, after 7 h FGF-1 treatment of astrocytes, we observed an increased single channel activity of Cx43 HCs. Although Cx43 HC activity was detected in spinal rat and mouse astrocytes after 7 h FGF-1 treatment, Cx43 HCs openings were rare in astrocytes maintained in control medium. These data indicate that the open probability of Cx43 HCs is increased after treatment of astrocytes with FGF-1; this increase is likely due to an increase in opening rate, since we did not detect an increase in surface Cx43 (Garré et al., 2010). Together these data provide molecular insight into how FGF-1 modulates the activity of Px1 HCs, Cx43 HCs, and P2X7Rs in spinal astrocytes.

In summary, the dual voltage clamp technique described here allows distinguishing the properties of different types of channels in astrocytes. By improving calculation of conductance in single channel recordings, we were able to discriminate the single channel activity of P2X7Rs and Px1 HCs, which cannot be achieved in astrocytes using conventional whole cell recordings. Also, this dual voltage clamp technique allowed us to compare activation of channels in the patches under strict control of V_m_. We anticipate that the dual voltage clamp system described here can be applied to human iPSC-derived glia and neurons to identify precise molecular and pharmacological therapeutic targets in neurodevelopmental and neurological disorders.

## Materials and Methods

### Ethics

Animal experimentation. Experimental procedures performed in making rat and mouse cell cultures were approved by the Institutional Animal Care and Use Committee of the Albert Einstein College of Medicine.

### Cell cultures

HeLa cells expressing Cx43-CFP or Px1-YFP were used as previously reported (Bukauskas et al., 2001; Garré et al., 2016). For one set of experiments, HeLa cells stably expressing Px1-YFP or Cx43-CFP and parental cells were seeded 1 day before experiments at low density. Single channel recordings were only conducted in isolated single cells.

For astrocyte cell cultures, rat spinal cords were removed from pups at postnatal day 1 (P1). Spinal cord tissue was mechanically dissociated and treated for 25 min with 0.001% trypsin-0.0002% EDTA solution in PBS (stock: 0.25% trypsin / 0.1% EDTA). Trypsin activity was stopped by adding 1 mM EDTA for 5 min, followed by further mechanical dissociation in maintenance medium composed of DMEM supplemented with 10% FBS, penicillin (100 UI/ml), and streptomycin (100 mg/ml). Cells were plated (at 2.0 × 10^4^ cells/cm2) on 12 mm glass coverslips or in 60 mm plastic culture dishes and kept in a maintenance medium. Cultures were grown at 37ºC in a humidified atmosphere of 5% CO2 and 95% O2.

For patch clamp experiments, only single isolated astrocytes were patched in sub-confluent monolayers 3 days after cell tissue isolation. Astrocyte cultures were > 90% pure as determined by their GFAP immunoreactivity, and there were no cells of myeloid linage (CD11b+ and CD11c+).

### Pannexin-1 and Cx43 deficient rat and mouse astrocytes

Subconfluent astrocyte cultures were treated with a sequence of Stealth siRNA duplex oligo ribonucleotides (siRNA Px1, 0.2 μM, ThermoFisher Scientific, catalog number 1330001) targeting rat Px1 mRNA and preventing Px1-mediated dye uptake (Garré et al., 2016). For control, we used a nontargeting siRNA, siRNA(−) (ThermoFisher Scientific). For transfection, confluent cultures (90%) were treated overnight with siRNA Px1 mixed with 10 μl/ml Lipofectamine reagent (ThermoFisher Scientific). Transfection reagents were removed and cells incubated for 4-6 h in DMEM medium (2% FBS) before 7 h treatment with FGF-1: Heparin (10 ng/ml: 5 IU/ml), as previously reported (Garré et al., 2016). Astrocytes were treated 7 h with heparin (50 IU/ml) as a control.

*Gja1*^*tm1Kdr*^ mice (Cx43 KO) (Reaume et al., 1995) were obtained from Dr. Eliana Scemes (Albert Einstein College of Medicine, Bronx, NY). Using the same protocol as that for rat astrocytes, Cx43^-/-^ astrocyte cultures were prepared from mouse embryos (E18), because pups die at birth of cardiac failure. The presence of targeted mutation was confirmed by PCR.

### Electrophysiology: dual voltage clamp

HeLa cells and astrocytes were grown on 22-mm number 0 coverslips and transferred to an experimental chamber mounted on the stage of an inverted microscope equipped with phase-contrast optics (IX70 microscope, Olympus). A fluorescence imaging system was used with HeLa cells to visualize Cx43 or Px1 with enhanced cyan fluorescent protein (Cx43-CFP) or yellow fluorescent protein (Px1-YFP) attached to the C-terminal.

Patch clamp recordings were conducted using the experimental chamber and either one or two EPC-8 patch clamp amplifiers (HEKA). The chamber was perfused with a modified Krebs-Ringer external solution containing (in mM): 140 NaCl, 4 KCl, 2 CaCl_2_, 1 MgCl_2_, 2 CsCl_2_, 1 BaCl_2_, 5 HEPES, 5 glucose, 2 pyruvate (pH 7.4). Patch pipettes for whole cell recording were filled with a solution containing (in mM): 10 NaAsp, 140 KCl, 0.2 CaCl_2_, 1 MgCl_2_, 2 MgATP, 5 HEPES (pH 7.3), 2 EGTA, 5 mM tetraethylammonium Cl-. Patch pipettes for cell attached recording were filled with the same solution as used for whole cell recording except that ATP was omitted to minimize activation of purinergic receptors. Since we generally opened the patch at the end of dual voltage clamp experiments, having both electrodes with very similar solutions should minimize changes associated with changing the internal solution. Pipette resistance was in the range of 5 to 10 MΩ. As a control for dual voltage clamp experiments, we performed experiments with external solution in the cell attached patch pipette, which revealed no differences from single channel conductances observed in symmetrical conditions. We also used external solution in all single clamp, cell attached patch experiments (**Figure S4**).

During a dual voltage clamp experiment, one pipette was in cell attached mode clamped to V_1_ = 0 to record channel activity (with a sensitivity of 50 mV/pA), and the other pipette (V_2_) was in whole cell mode to clamp V_m_ = V_2_ - V_1_ and stimulate (recording sensitivity of 1mV/pA). In some experiments to validate the method, the gain in the amplifier connected to the whole cell pipette was increased to 50 mV/pA to simultaneously record channel activity at the same gain with both pipettes.

Voltages and currents were recorded and subsequently digitized using a MIO-16X A/D converter and acquisition software (Bukauskas et al., 2001). Records were digitized at a rate of 5 kHz and then passed through a digital Gaussian filter with pass frequency between 0.2-0.5 kHz.

### Single channel analysis

Channel gating events were identified as rapid transitions, opening and closing, between periods at constant voltage (steps) or during the linearly changing phase of voltage ramps. Repeated transitions of fixed amplitude (or conductance) from the basal level indicated activity of a single channel. Single channel events were characterized by three parameters, membrane potential, current change between open and closed states, and duration. One cell attached patch per cell was established with a seal resistance of 1-5 GΩ from 3-15 cells for each experimental condition. For dual voltage clamp experiments, single-channel conductance was calculated from single channel currents in the cell attached pipette divided by the difference between the voltage in the whole cell and cell attached recordings (V_2_-V_1_). We defined V_1_ as the electrode initially used in cell attached mode, and shifted to whole cell mode at the end of the experiment, whereas, V_2_ was the electrode used in whole cell mode during the entire period of recording. A correction factor, V_1_/V_2_, was calculated at the end of each experiment by opening the attached patch, shifting amplifier mode to current clamp in both pipettes and passing currents through V_1_ comparable to those used during the experiment. V_1_/V_2_ was generally close to 0.8-1 at the end of the experiment, indicating that the bridge circuit that removes the voltage drop in the whole cell pipette was properly balanced. To present currents in the same direction in both whole cell and cell attached patch recordings, actual currents recorded through the V_1_ electrode were multiplied by -1.

### Statistics

Data are presented as dot plot distributions and mean ± SEM; n expresses the number of ramps, trials, or patches per cell from a minimum of 3 independent experiments (*i*.*e*., HeLa cells split on different days; astrocyte cell cultures prepared from ∼10 pup spinal cords from at least 3 different litters). Means for each group were compared using a non-parametric test for continuous and non-paired variables, the Mann-Whitney test or paired Wilcoxon test; and one-way analysis of variance (ANOVA). Statistics were performed using Graph Pad Prism 9 (2021) software. Differences were considered significant at *P* < 0.05. Graphics were prepared using Adobe Illustrator (2021), Sigma Plot (2000) and Graph Pad Prism 9.3 (2021).

## Acknowledgments

This work was supported by National Institutes of Health Grants NS45287 and NS55363 to M.V.L.B. and NS072238 to F.F.B.

