## Supplemental Figures for "A dual voltage clamp technique to study gap junction hemichannels in astrocytes cultured from neonatal rodent spinal cords"

**Supplementary figures**

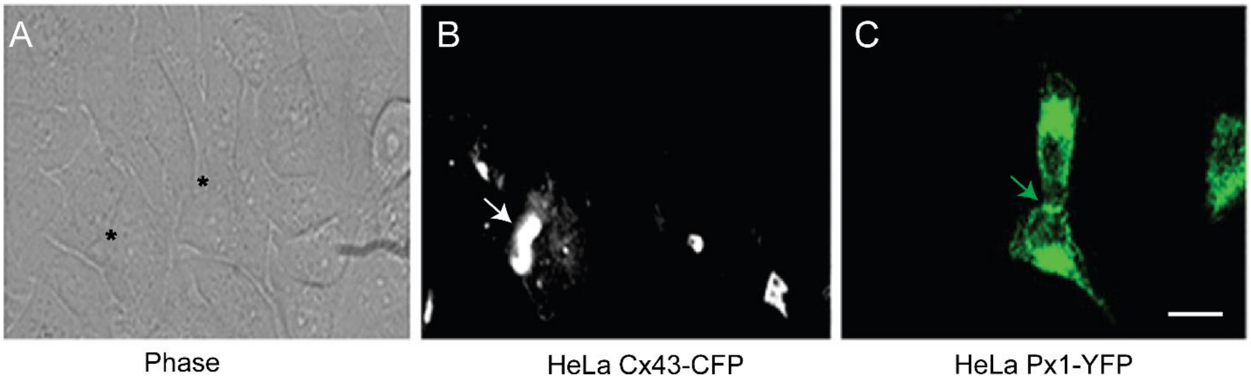


**Figure S1**. Representative images of mixed cultures of HeLa cells overexpressing rat Cx43-CFP (gray) and human Px1-YFP (green). Both Cx43 and Px1 were detected in putative sites of gap junction formation, intracellularly, and in the unopposed cell membrane. (**A**) Phase image, asterisks indicate cells expressing Cx43-CFP and Px1-YFP. (**B**) Plaque-like labeling is present between HeLa Cx43-CFP cells (white arrow), indicating putative sites of Cx43 HC intercellular docking and Cx43 GJ channel formation. (**C**) In contrast, plaques like those in HeLa Cx43-CFP cells were not observed between HeLa Px1-YFP cells, although a small labeled plaque (green arrow) between cells suggests formation of Px1 GJ channels. Scale bar 10 μm.

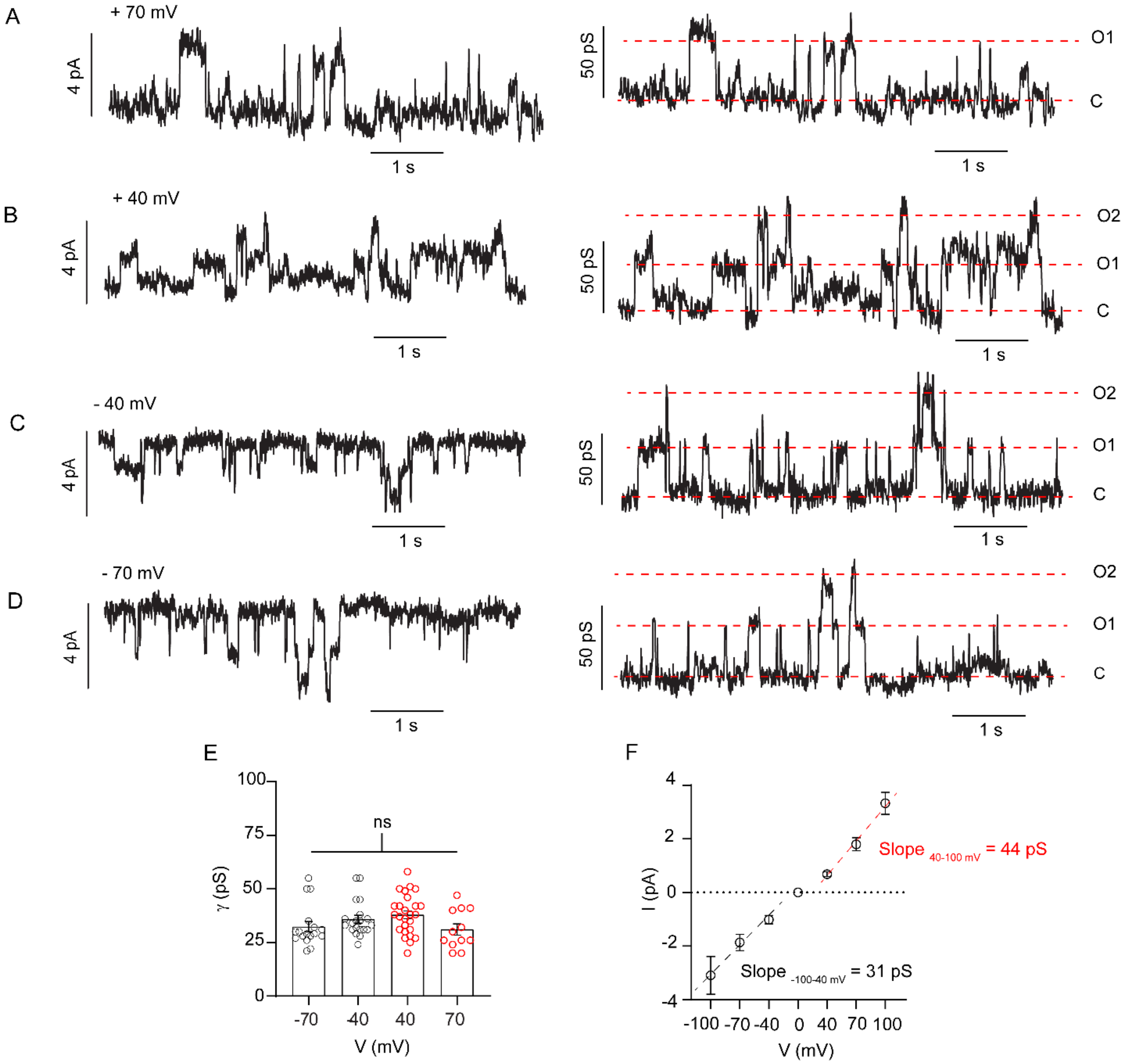


**Figure S2**. Single channels recorded with whole cell patch clamp in HeLa Px1-YFP cells. (**A-D**) Whole cell currents (left panels) and the respective single channel conductances (right panels) at 70, 40, -40, -70 mV. (**E**) Single channels opened with little rectification and unitary conductances of 37.9 ± 9.3, 31.1 ± 9.1, 32.4 ± 9.8, 35.9 ± 8.2 pS at -70,- 40, 40, 70 mV, respectively ($n$ = 17, 20, 26, 12, recordings, one whole cell recording per cell, ns, not significant, one-way ANOVA). (F)

Single channel I-V curves and calculated slopes, 31 pS and 44 pS, at negative (-100 to 40 mV, black) and positive (40-100 mV, red) $V_m$, respectively.

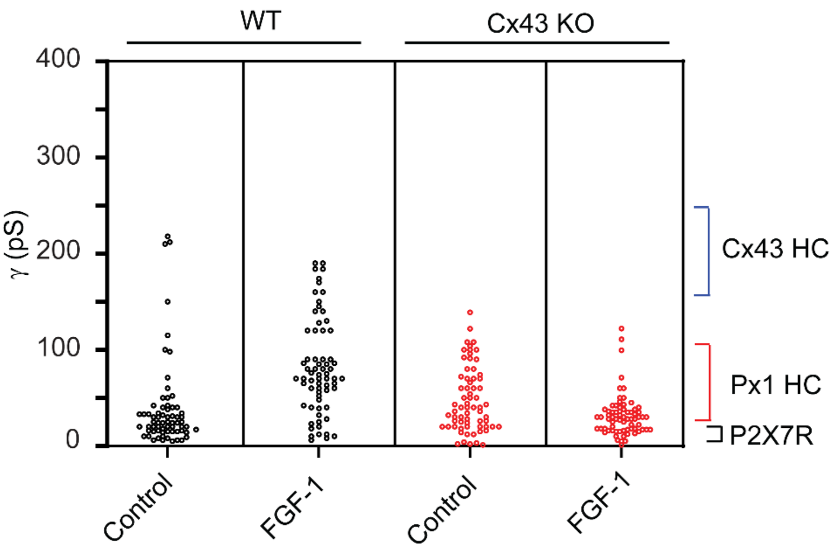


**Figure S3.** Dot plots for visualizing the distributions of conductance of single channel openings recorded in cell attached patches in mouse spinal astrocytes. Responses were evoked by ramps of 1.7 s duration, -/+ 80 mV amplitude through a whole cell pipette in spinal astrocytes prepared from *WT* and Cx43 KO embryos. Cells were maintained in control medium or in medium supplemented with FGF-1 for 7 h ($n$ = 70 ramps, 5 ramps per patch, 14 patches, one patch per cell, for each condition). Similar to rat astrocytes, the amplitude and frequency of unitary events were increased after treating mouse astrocytes for 7 h with FGF-1. Cx43 HC openings were not detected in Cx43-/- astrocytes control and treated with 7h FGF-1.

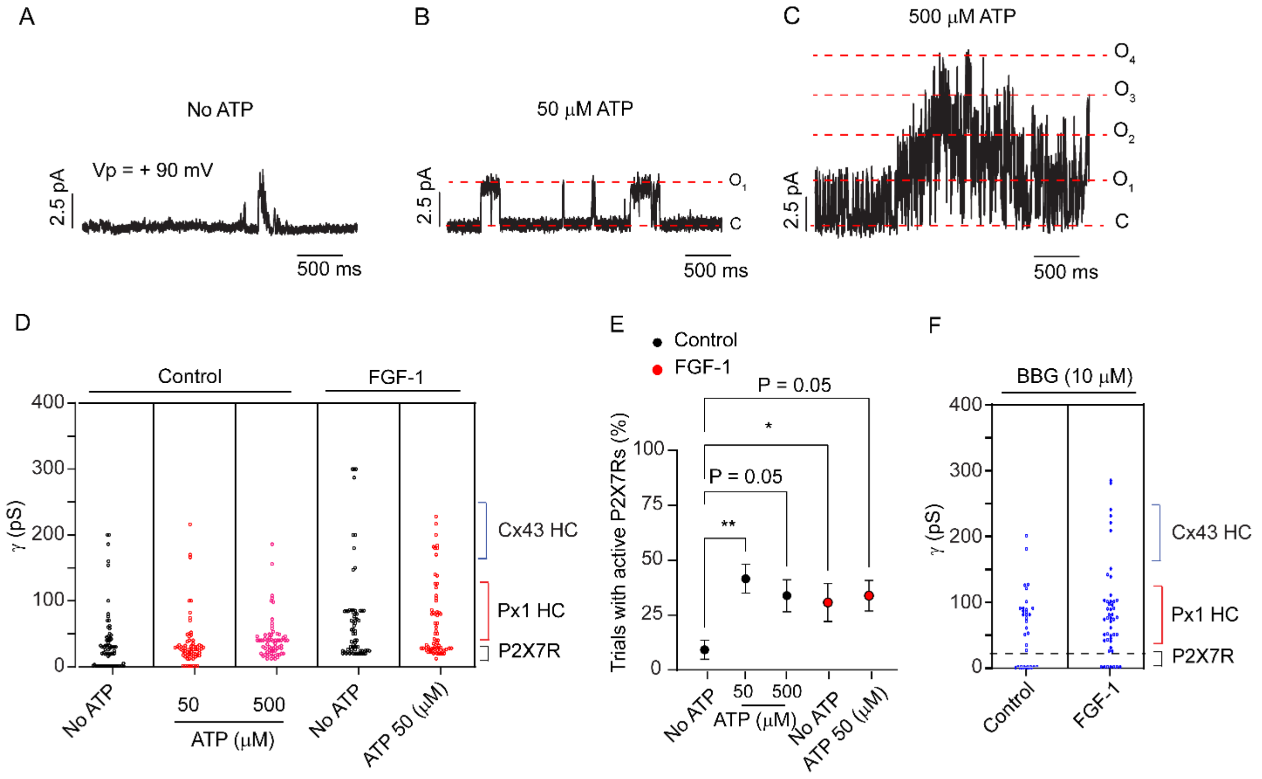


**Figure S4**. ATP-induced unitary currents in spinal astrocytes cultured from neonatal rat spinal cord. (**A-D**) Unitary events were recorded using cell attached patch single clamp. Voltage in the cell attached pipette ($V_p$) was + 90 mV so that voltage across the patch was ~110 mV positive to the resting potential. The pipette was filled with normal external solution. ATP was not added to the pipette solution (**A**), or added at concentrations of 50 µM (**B**) and 500 µM (**C**). (**D**) Dot plot distributions of single channel transitions (both opening and closing) in astrocytes maintained in control medium (with no ATP in the pipette, or with 50 or 500 µM ATP in the pipette) and in astrocytes treated with 7 h FGF-1 (with no or with 50 µM ATP in the pipette). (**E**) Percentage of trials (voltage pulses) ($V_p$ held during ~10 s at + 90 mV, relative to $V_{m,}$) with active P2X7Rs when ATP was not added to the pipette solution, or added at concentrations of 50 µM and 500 µM in control or 7h FGF-1 treated spinal astrocytes ( *n* = 12 trials, 2 trials/patch, from 6 different patches and cells). *$P$ < 0.05, **$P$ < 0.01, ns, not significant, one-way ANOVA followed by Sidak's multiple comparisons test). (**F**) BBG prevents P2X7R openings, but it does not affect single channel activity of Px1 HCs or Cx43 HCs. Dot plot distributions from cell attached patch clamp recordings in control and FGF-1-treated astrocytes in which BBG (10 μM) and ATP (50 μM) were in the cell attached patch electrode. Events were analyzed from 69 trials with $V_p$ held at + 90 mV, relative to $V_m$ ( $n$ = 7 patches, ~10 trials/patch, one patch per cell).
